# Fecal Microbiota Transplantation in a Domestic Ferret Suffering from Chronic Diarrhea and Maldigestion–Fecal Microbiota and Clinical Outcome: A Case Report

**DOI:** 10.1101/2023.11.20.567889

**Authors:** Sean J. Ravel, Victoria M. Hollifield

## Abstract

This case report describes the effects of fecal microbiota transplantation (FMT) administered via enema in a 4-year-old spayed, champagne Domestic Ferret (*Mustela putorius furo*) with chronic diarrhea, maldigestion and weight loss. We aimed to establish a protocol for FMT as a novel therapeutic treatment for chronic diarrhea in domestic ferrets. We mapped the fecal microbiome by 16S rRNA gene amplicon sequencing to track the patient’s fecal microbiota throughout the treatment and observation period. Initial oral FMTs were associated with a temporary weight improvement but subsequent treatments, via enema and oral delivery, showed varied outcomes. Molecular analysis highlighted distinct gut microbiota composition profiles between the healthy donor and the diseased ferret. The diseased ferret initially exhibited high abundance of *Enterobacteriaceae, Escherichia*, and *Enterobacter*, which ultimately normalized to level like those found in the donor ferret. Overall, the gut microbiota of the recipient became more similar to the donor microbiota using a Yue-Clayton theta coefficients analysis. After a restoration of the gut microbiota and clinical improvement, the recipient’s symptoms returned. Future studies are warranted to map the microbiome of a larger population of domestic ferrets to investigate a potential correlation between fecal microbiota profiles and chronic/acute gastrointestinal disorders.

## Introduction

In recent years, Fecal Microbiota Transplantation (FMT) has shown promise for treating dysbiosis-related gastrointestinal disorders in human and animal models^1,2^. The procedure consists of grafting feces from a healthy donor into the gut of a diseased recipient to establish eubiosis in the gut of the recipient. In humans, the procedure has been successfully used to treat *Clostridioides difficile* infections^3^ and specific products have been approved by the FDA for treating this devastating disease^4-6^. In animals, while FMT has been used for many years to treat indigestion in ruminants^7,8^, the use of FMT is rarely used by small animal practitioners^8,9^. The gut microbiota has been documented to play an essential role in maintaining the overall health of humans, dogs, cats, and other species by providing protection from transient enteropathogens, regulating the immune system, supplying metabolites necessary for proper nutrition, and other vital functions^10-13^.

No case reports have been published in the literature on the use of FMT in the domestic ferret. Ferrets are increasingly popular companion animals as well as a valuable animal model for human vaccine development and research of various human gastrointestinal infections such as campylobacteriosis, *C. difficile* and *Helicobacter pylori* gastritis^14-16^. Based on the success in other animals^17,18^, we investigated the potential clinical use of FMT to treat ferrets in the research and companion pet industries.

In this case report, we describe the use of FMT in the case of a ferret suffering from chronic diarrhea, being unresponsive to multiple antibiotics, and anti-diarrheal therapies. We aimed to establish an FMT protocol and assess the effectiveness of FMT as a novel therapy for chronic diarrhea in ferrets.

## Clinical Case Description

### Patient History and Clinical Presentation

The case was an approximately 4-year-old spayed Domestic Ferret, surrendered to Best Friends’ Veterinary Hospital (Gaithersburg, MD, USA) in September of 2022, presenting with chronic diarrhea, adrenal gland disease, and suspected insulinoma. The patient was administered a subcutaneous deslorelin acetate (4.7mg; Suprelorin^®^ F, Virbac, Carros, France) implant in October 2022 and prescribed a continuous low prednisolone dose (0.8mg per os SID; Lloyd, Inc., IO, USA) to manage the adrenal gland disease and insulinoma, respectively. The diarrhea was characterized by very loose, seedy, and mucoid stools, indicating that food passed through the patient’s GI tract too quickly.

The patient was transitioned to a commercial ferret diet for digestive support (Ferret Wysong Epigen 90™ Digestive Support, Wysong, MI, USA). This diet was supplemented twice daily with ¼ teaspoons of pancreatic enzyme powder (Pancreazyme^®^, Virbac AH, Inc., TX, USA), 0.4mL solution (1:30) of milk thistle seed extract (33.4mg/mL; Nature’s Answer, NY, USA) and lactulose (0.67mg/mL) (Hi-Tech Pharmaceuticals, GA, USA), and 1mL aqueous suspension of activated attapulgite (150mg/mL, Ensdosorb^®^, PRN Pharmacal, FL, USA). Additionally, 2-3g of a high-calorie vitamin supplement (FerretVite^®^, Spectrum Brands, Inc., VA, USA) and 1g of probiotic gel (Bene-bac Plus^®^, PetAg, Inc., IL, USA) were administered three times daily. The patient chronic diarrhea persisted after several weeks; however, appetite and water intake were noted to be normal and regular.

In November 2022, the patient was prescribed a regimen of bismuth subsalicylate (0.8mL of 7.9mg/mL *per os* BID q7d; PeptoBismol^®^, Procter & Gamble, OH, USA), metronidazole suspension (0.45mL of 16.67mg/mL *per os* BID q21d; A.P.I., Inc., GA, USA), amoxicillin suspension (0.5mL of 50mg/mL *per os* TID q21d; Sandoz, Inc., NJ, USA), and two bi-daily probiotic supplements (Bene-bac Plus^®^ & ProViable®-DC Capsules, Nutramax Laboratories Veterinary Sciences, Inc., SC, USA). After the complete 21-day treatment regimen, diarrhea remained unresolved. The patient’s body weight dropped 5% (800g to 760g) 30 days post-treatment. Fecal gram-stain tests conducted post-treatment showed extremely low bacterial abundance, which was cause for serious concern. At this stage, oral FMT was considered for this patient as a novel therapeutic treatment for chronic diarrhea.

Over four days following the administration of the oral FMT, the patient’s body weight increased from 763g to 801g before dropping to 762g on the 5^th^ day. The stool consistency increased to 3/5 for the four days post-procedure before dropping to 1/5. No samples for microbiota characterization were collected over this initial treatment period. After showing promises at reducing the patient’s diarrhea and weight loss, the oral method of administration was originally abandoned for the enema under the presumption that colonization would be more efficient under the hypothesis that direct deposition of the fecal slurry to the large intestine would allow for microorganisms better colonize the gut environment. The study protocol and associated fecal microbiota characterization described below started 7 days after this initial oral FMT.

## Materials and Methods

### Donor Candidate Selection and Characteristics

Two candidates were considered using a screening protocol criteria that included age (<4 years); regular core vaccinations; history of high-quality diet; no history of acute/chronic gastrointestinal diseases; no antimicrobial administration; satisfactory body condition score (4-5 on a 9 point scale); normal fecal consistency; no history of chronic disorders (immune-mediated, endocrine, and neoplastic); a negative centrifugal fecal flotation with zinc sulfate solution test for helminth ova; and a satisfactory clinical physical examination.

The donor selected in this case was a healthy 3-year-old, 1.4 kg male neutered domestic ferret with a body condition score (BCS) of 5/9 with no history of gastrointestinal disease or chronic disorders. The donor was currently vaccinated and dewormed. Additionally, there was no history of antibiotics use, but they were presumably prescribed post-castration in 2019 before the owner obtained the ferret.

The donor ferret was fed the same diet as the recipient ferret (Wysong Epigen 90™ Digestive Support) and was not taking any medications. Fecal consistency had been normal, with no diarrhea in the last +24 months. Fecal floatation was negative for helminth ova, *Giardia* spp., and other protozoa. Two fecal gram-stains seven days apart were 100% gram-positive, and no *Campylobacter* spp., *Clostridia* spp., or yeast were observed.

### Preparation of Donor Sample

This protocol is derived from the protocol developed by Furmanski et al. established for cats with the following modifications^18^. The donor ferret was kept in a separate cage at the veterinary hospital, and a freshly voided sample was collected within 30 minutes. It was immediately weighed and a 0.17mg/mL slurry generated in warmed (∼30ºC) and sterile 0.9% NaCl. The sample was filtered three times through two sterile gauzes soaked in warmed and sterile 0.9% NaCl to remove large particulate matter and drained into a sterile container. The fecal preparation was transferred within 30 minutes to the recipient. The procedure was repeated for each transplant performed.

### Preparation of the Recipient for FMT by Enema

Ferrets have a relatively short digestive transit time of 3-4h compared to dogs and cats, 6-8h and 10-20h, respectively^19^. The patient was fasted for 3 hours before and 1 hour after the procedure to prevent premature evacuation of the bowels. Prior to the enema FMT (eFMT), the ferret was anesthetized using isoflurane according to standard clinical procedures.

### Administration of FMT by Enema

The required volume (10mL/kg body weight) of 8mL was drawn up in a 10 mL syringe with a FR 6.5 330mm Foley Catheter. The catheter was then prefilled with fecal slurry to prevent introducing air to the colon. The catheter was lubricated using veterinary lubricant jelly and gently advanced up to the estimated location of the transverse colon while the anus was held closed by a technician. After the 8 mL of fecal slurry were deposited in the patient’s colon, it was taken off anesthesia and held with its caudal body slightly elevated for approximately 5-10 minutes while recovering. One-hour post-procedure, the patient was given its regular diet and medications. The procedure was performed on day 1 and repeated on day 3 (**Figure 1**).

**Figure 1.**
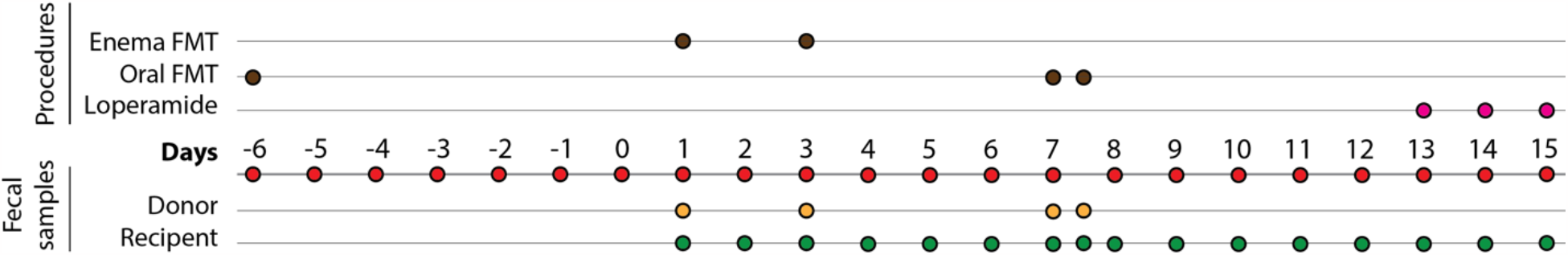
Timeline. FMT and sample collection.

### Oral FMT Maintenance Doses

To support gut microbiota colonization, a maintenance dose was administered *per os* twice on day 7 (morning and evening, 10 hours apart) (**Figure 1**). A fecal slurry was prepared following the same protocol described above but with an increased concentration (0.25mg/mL of stool in warmed (45ºC) and sterile 0.9% NaCl). The patient was fasted for 1 hour before the oral FMT (oFMT) procedure to prevent overfilling the stomach. A technician scruffed the ferret, and 2 mL of fecal slurry was deposited into the ferret’s mouth at a rate of 2 mL per minute with a 3 mL oral syringe. The oFMT was immediately followed with 2-3 grams of a malt syrup-based vitamin supplement (FerretVite) to mitigate discomfort for the patient and prevent regurgitation.

### Fecal Sample Collection

Fecal samples from the recipient were collected immediately after voiding on days 1 through 15. Samples of the donor feces (1 g) and 1mL fecal slurry used in each procedure were collected and analyzed. Additionally, control environmental samples (air) were periodically collected from the area inside and around the patient’s cage throughout the study. All samples were placed in 1mL of nucleic acid stabilizing solution (DNA/RNA Shield, Zymo Research) in a 2 ml Hamilton FluidX tube and stored at -80°C until processing for DNA extraction and 16S rRNA gene amplicon sequencing.

### 16S rRNA Gene Amplification and Amplicon Sequencing

DNA extraction was performed from 200μL of fecal sample suspension in DNA/RNA Shield using the MagAttract Microbiome DNA kit (QIAGEN) automated on a Hamilton STAR robotic platform and following the manufacturer’s instructions. Amplification and amplicon sequencing of the V3-V4 regions of the 16S rRNA gene was performed according to a validated and previously published protocols and sequenced on an Illumina MiSeq instrument^20^ using the PE300 protocol by the Maryland Genomics Core at the Institute for Genome Sciences, University of Maryland School of Medicine, Baltimore, Maryland, USA. Controls samples included an 8-member positive controls (Zymo Research), as well as an extraction negative and a PCR negative controls. Sequence data processing, quality controls and taxonomic assignment were performed using the bioinformatic pipeline developed by Holm *et al*. ^20^ resulting in a table of bacterial phylotypes relative abundances. Taxonomic assignments were performed using the SILVA database release 138.1^21^. For the analysis, controls and phylotypes present at less than 10^−4^ study wide were removed from the datasets. This resulted in a taxonomic table comprising 31 unique phylotypes (Supplementary Table 1). All analyses and graphs were performed in R version 4.3.1.

A total of 3,402,051 16S rRNA gene amplicon sequences were obtained from 23 stool samples (mean number of sequences per samples was 156,723 (SD: 49,030, range: 2,184 to 211,482). Environmental controls did not generate any sequences.

### Clinical Observation

Following the first eFMT, clinical symptoms were recorded daily including attitude/activity; stool consistency; and body weight. Stool consistency was ranked on a 5-point scale with 1 being extremely liquid and 5 being healthy and well-formed stool. Attitude/activity were assessed subjectively and ranked on a 5-point scale with 1 being nearly catatonic and 5 being extremely hyperactive. Observations were recorded daily.

## Results and Discussion

### Clinical Outcomes

Clinical observation data are summarized on **Figure 2**. The initial oFMT led to a major improvement of the patient’s weight which increased from 763g to 801g in 4 days (day -2 on **Figure 2**).

**Figure 2.**
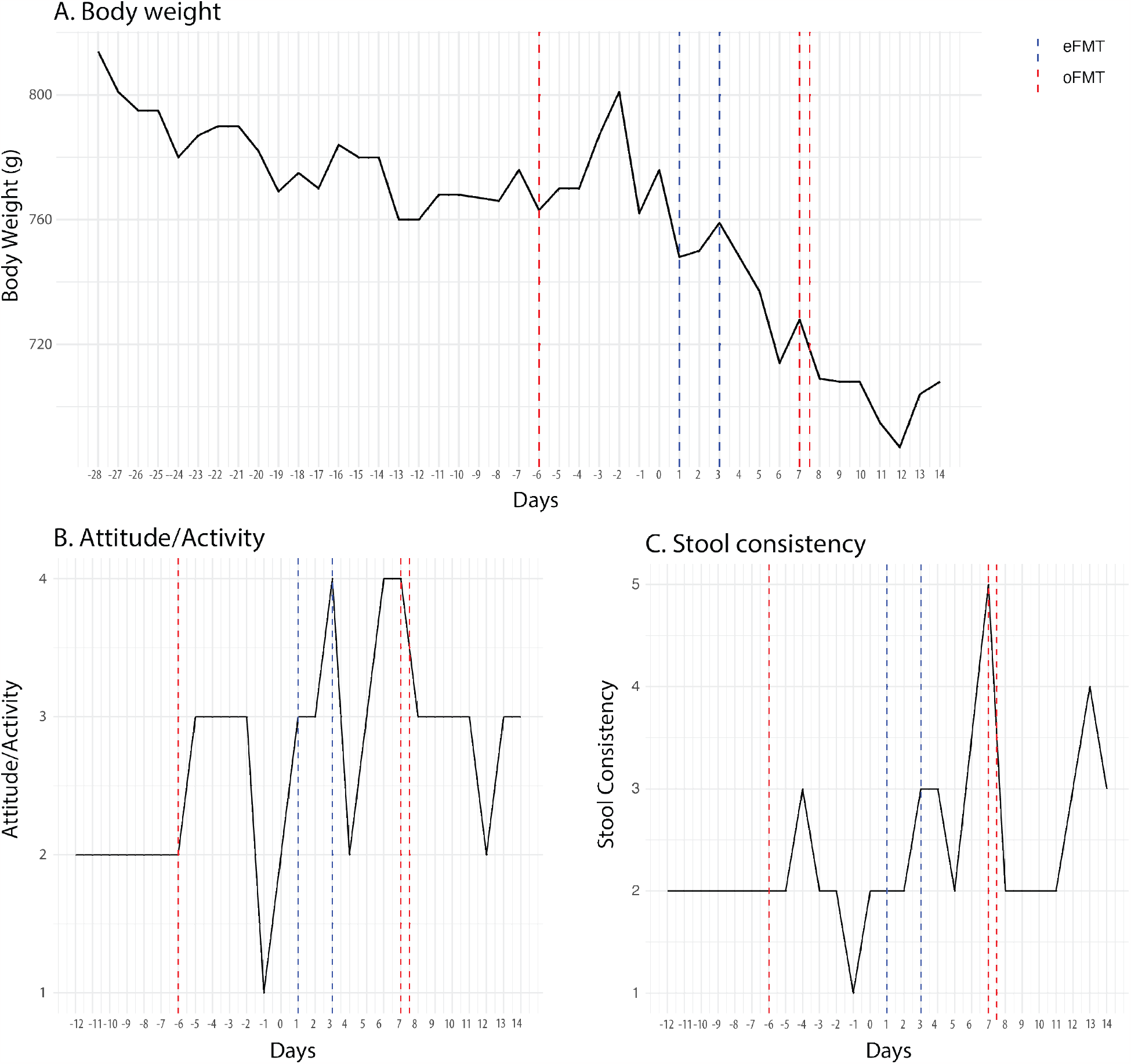
Clinical observations during the duration of the study including body weight (A), daily attitude/activity (B), and daily stool consistency (C). Enema FMT (blue) and oral FMT (red) administrations are represented by vertical bars.

Unfortunately, the effect was not sustained, and the patient’s weight dropped back to 762g five days after the procedure. At that time, it was decided to perform an eFMT to maximize delivery of the fecal slurry to the gut. Samples were collected for microbiota analysis before and after the eFMT. Two eFMT were performed on day 1 and day 3 using two separate stool samples from the same donor collected within 30 minutes of the procedure. Unlike the initial oFMT, no improvement in the patient’s body weight was observed. The patient’s weight dropped from 748g on day 1 to 737g on day 5. Similarly, no improvement in stool consistency and attitude/activity levels were observed, and both remained low. Gram-stain examination of a day 5 fecal sample revealed very low bacterial abundance. On day 6, the patient’s stool consistency increased to 3.5; the best consistency observed since observations began in September 2022, as well as its attitude and activity levels which increased drastically. However, its body weight decreased to 714g. On day 7, the patient’s body weight increased to 728g, and stool consistency improved to 5/5. On that day, two FMTs by oral gavage were performed 12 hours apart. Unfortunately, over the next 5 days, the patient’s body weight declined to 687g at which point the patient was started on a loperamide hydrochloride solution (0.93mL of 0.13mg/mL *per os* TID) and daily subcutaneous fluids (8-10 mL 50:50 NaCl/Lactated Ringer w/ 5% Dextrose SID). The rest of the planned oFMTs were abandoned for the safety of the patient. After starting the loperamide, the stool became less mucoid and more formed; however, the seedy texture remained indicating persistent maldigestion.

Interestingly, Gram-stains were conducted periodically after the last oFMT on day 6 up to day 15. These tests revealed a gradual increase in bacterial abundance up to day 15 at which point it was subjectively judged to be normal. All tests were observed to be 95-100% gram-positive bacteria.

### Fecal Microbiota Analysis

We performed a molecular analysis of the composition of fecal microbiota in the donor and recipient. We further aimed to evaluate if the recipient fecal microbiota composition was converting to that of the donor. The gut microbiota of the healthy donor ferret was primarily characterized by the predominant presence of *Clostridium* spp. While *Clostridium* spp. were the major taxa, other bacterial taxa were observed albeit at lower abundance, including *Turicibacter, Peptostreptococcaceae*, and *Romboutsia* (**Figure 3**). Interestingly, these never colonized the gut of the diseased ferret at level similar to that of the donor. The composition of the fecal microbiota of the donor ferret appears to be quite dynamics with fluctuating relative abundance of different bacterial taxa (**Figure 3A** and **3B**).

**Figure 3.**
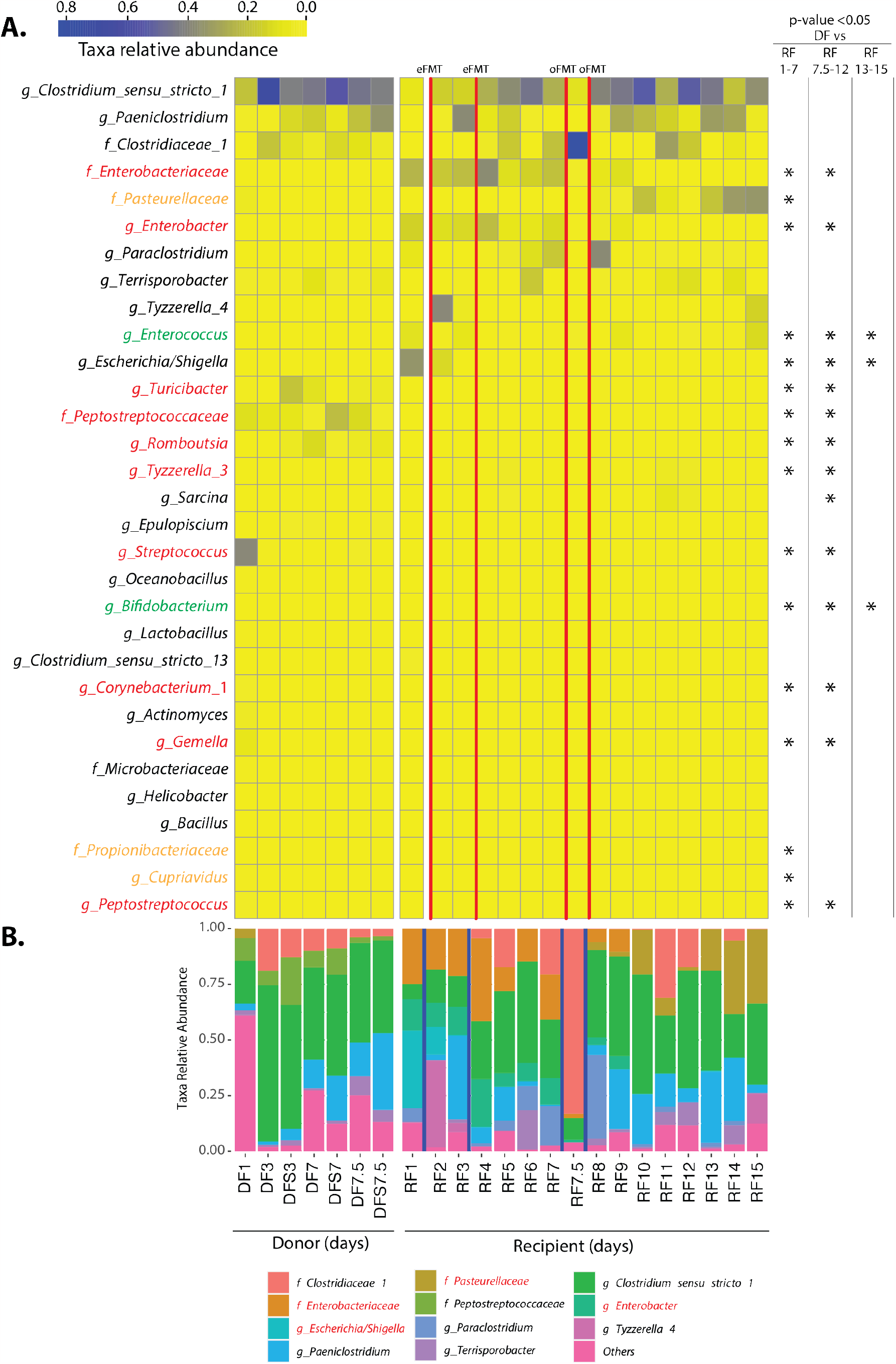
A. Heatmap showing the relative abundances of all 31 bacterial taxa detected in the fecal microbiota of the donor (DF) and recipient (RF) fecal samples. The stars indicate taxa statistically equally abundant over between all the donor samples combined and the recipient over the period indicated (day 1-7: after eFMT, day 7.5-12: after oFMT, and day 13-15: after initiation of loperamide) (statistical test was Kolmogorov-Smirnoff test, p-value < 0.05). Taxa in green were similar at all three period, taxa in red were similar at the first two periods, while taxa in orange were similar only at the period after eFMT. **B**. Stacked bar plot showing the relative abundance of the top 11 bacterial taxa for each sample. The number associated with the donor (DF and DFS) and recipient (RF) indicates the day of collection. Note that two separate samples were collected on day 7. DF: Donor fecal sample, DFS: Donor transferred fecal slurry; RF: recipient fecal sample.

The microbiota of the diseased ferret was markedly different, characterized by a dominance of *Enterobacteriaceae* during the first 7 days, with a high abundance of *Escherichia/Shigella* prior to the eFMT on day 1. While *Enterobacteriaceae* decreased throughout the treatment period, *Pasteurellaceae* increased after the two oFMT on day 7, to reach relative abundances of more than 15% on day 13. Interestingly, *Escherichia/Shigella* rapidly diminished following the eFMTs and were undetectable for the remainder of the sampling period.

To evaluate the over similarity of the microbiota between the donor and the recipient, we calculated the Yue-Clayton theta coefficients between each recipient samples and the average of the donor samples (**Figure 4)**. On the first day, the Yue-Clayton theta coefficient was 0.868, indicating high dissimilarity. Following the administration of the first eFMT, the coefficient decreased to 0.702 on day 2 and continued decreasing after the second eFMT on day 3, reaching a pronounced low (high similarity) of 0.138 on the day 5. However, on the seventh day, the coefficient sharply increased to 0.726. After the two oFMT, the coefficient decreased and reached a value of 0.072 on the day 12. The remaining days of the study saw fluctuations in the coefficient, trending upwards.

**Figure 4.**
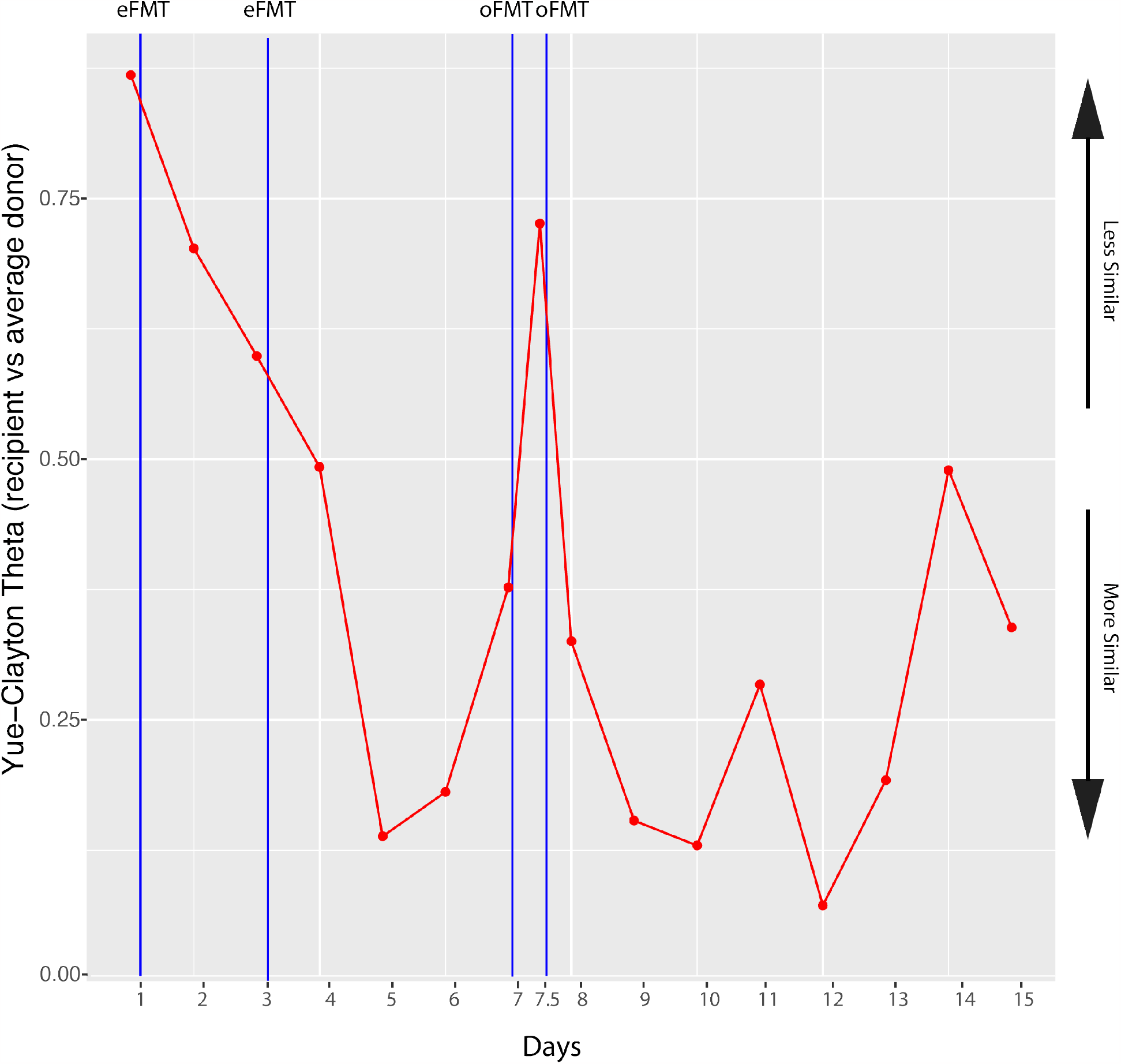
Comparison of the composition of the fecal microbiota of the donor to the recipient. Yue-Clayton theta values are plotted for each recipient fecal microbiota compared to the average composition of the donor fecal microbiota. The lower Yue-Clayton theta value is the more similar are the microbiota composition.

## Conclusions

We aimed to use FMT to reestablish eubiosis of the gut microbiome in a ferret experiencing severe diarrhea and weight loss and restore stool consistency. Gram-stain analysis of the patient’s stool demonstrated a low bacterial abundance and severe dysbiosis, which were likely contributing factors to the observed chronic mucoid, seedy diarrhea and body weight loss.

Microbiota composition evaluation pointed to the presence of *Enterobacteriaceae* and *Enterobacter* as the likely cause of the condition, as well as *Escherichia*. The administration of FMT by enema, followed by oral FMT as administered in this study only ameliorated the patient symptoms temporarily, lowered the amount of *Enterobacteriaceae* but ultimately, did not alleviate the symptoms and treatment had to be stopped, possibly indicating that the origin of the symptoms be only in part associated with a dysbiotic gut microbiota. The patient’s co-morbidities (suspected insulinoma, adrenal gland disease) may have contributed to the limited success observed.

This is the first report of the use of FMT to treat severe diarrhea in ferret. While the procedure was not successful in this study, further research is needed improve the procedure and positively modulate the composition of the ferret gut microbiota. Unfortunately, little is known about the composition of the ferret gut microbiota and research aimed to map the ferret fecal microbiota and its association with health and disease is warranted. This research would contribute greatly to advancing the potential of this species as a research model as well as adding to the existing clinical veterinary medicine knowledge for agricultural and companion pet care.

## Supporting information

Supplementary Table 1 Taxonomic Table

## Acknowledgements

The authors would like to extend their deepest gratitude to the staff of Best Friends’ Veterinary Hospital for the outstanding patient care they provided to the patient and donor ferret.

## Author Contributions

SJR conceptualized the study, conducted the primary data collection, analysis, and interpretation, and led the writing of the manuscript. VMH contributed to the experimental design and revision of the manuscript. Both authors, SJR and VMH, have read and agreed to the published version of this manuscript.

## Ethic Statement

The clinical procedures and sample collection were performed as part of regular practice of veterinary medicine under the supervision of Victoria M. Hollifield, DVM. In this instance, informed consent for the treatment and subsequent publication of the case report was not required as the ferret had been surrendered to the hospital, granting Best Friends’ Veterinary Hospital ownership and the right to make medical decisions in the animal’s best interest. This case report was prepared with the highest regard for the ethical treatment of animal subjects. All procedures were performed by qualified individuals and designed to minimize distress and discomfort. This report reflects a commitment to ethical scientific inquiry and respect for the lives of animal subjects.

## Supplementary Materials

**Supplementary Table 1:** Microbiota taxonomic table of donor and recipient samples

